# L1PA2 transposons contribute abundant regulatory sequences in MCF7 breast cancer cell line

**DOI:** 10.1101/2020.09.01.276808

**Authors:** Jiayue-Clara Jiang, Joseph Rothnagel, Kyle Upton

**Affiliations:** School of Chemistry and Molecular Biosciences, The University of Queensland, St Lucia, QLD 4072, Australia

**Keywords:** Transposons, L1PA2, breast cancer, transcription factor binding sites, transcription start sites

## Abstract

Transposons, a type of repetitive DNA elements, can contribute cis-regulatory sequences and regulate the expression of human genes. L1PA2 is a hominoid-specific subfamily of LINE1 transposons, with approximately 4,940 copies in the human genome. Individual transposons have been demonstrated to contribute specific biological functions, such as cancer-specific alternate promoter activity for the *MET* oncogene, which is correlated with enhanced malignancy and poor prognosis in cancer. Given the sequence similarity between L1PA2 elements, we hypothesise that transposons within the L1PA2 subfamily likely have a common regulatory potential and may provide a mechanism for global genome regulation. Here we show that in breast cancer, the regulatory potential of L1PA2 is not limited to single transposons, but is common within the subfamily. We demonstrate that the L1PA2 subfamily is an abundant reservoir of transcription factor binding sites, the majority of which cluster in the LINE1 5’UTR. In MCF7 breast cancer cells, over 27% of L1PA2 transposons harbour binding sites of functionally interacting, cancer-associated transcription factors. The ubiquitous and replicative nature of L1PA2 makes them an exemplary vector to disperse co-localised transcription factor binding sites, facilitating the co-ordinated regulation of genes. In MCF7 cells, L1PA2 transposons also supply transcription start sites to up-regulated transcripts. These transcriptionally active L1PA2 transposons display a cancer-specific active epigenetic profile, and likely play an oncogenic role in breast cancer aetiology. Overall, we show that the L1PA2 subfamily contributes abundant regulatory sequences in breast cancer cells, and likely plays a global role in modulating the tumorigenic state in breast cancer.

## INTRODUCTION

Transposons are repetitive DNA elements that are ubiquitous in eukaryotic genomes, and occupy about 45% of the human genome (Lander et al. 2001). Transposons contain cis-regulatory sequences that are recognised by the host transcriptional machinery (primarily transcription factors (TF) and RNA polymerases (RNA pol)), which they require to exploit the host transcriptional resources for their own replication (Jordan et al. 2003; Cruickshanks and Tufarelli 2009; Gifford et al. 2013; Sundaram et al. 2014; Chuong et al. 2016; Chuong et al. 2017). This obligatory compatibility also enables the host genome to exapt transposon-derived regulatory sequences to modulate the regulation of host transcriptional networks.

Transposons have been shown to contribute functional sequences, including TF binding sites (TFBS), promoters, enhancers and insulators, for the regulation of host genes in a variety of organisms, including humans, mice, maize and fruit flies (McClintock 1956; Gerlo et al. 2006; Flemr et al. 2013; Guio et al. 2014; Notwell et al. 2015; Chuong et al. 2016; Ding et al. 2016). Host genes that are under the regulation of transposon sequences are involved in a diverse range of biological functions, including innate immunity and pregnancy in humans, female fertility and brain development in mice, as well as courtship song phenotype and stress response in fruit flies (Gerlo et al. 2006; Flemr et al. 2013; Guio et al. 2014; Notwell et al. 2015; Chuong et al. 2016; Ding et al. 2016).

In addition to gene regulation in normal biological pathways, transposons may also play a regulatory role in the cancer transcriptome. In the human genome, transposons are often heavily suppressed in somatic tissues, primarily via histone tail modifications and DNA methylation; however, they can escape epigenetic repression in the cancer state, and exert regulatory activity that affects the expression of oncogenes (Choi et al. 2009; Sharma et al. 2010; Lee et al. 2012; Babaian and Mager 2016). The transposon-derived activation of oncogenes, called the onco-exaptation of transposons, can subsequently lead to tumour development and enhanced malignancy, as summarised in Babaian and Mager (2016). Specific examples of transposon-driven abnormal oncogene expression have been reported in human lymphoma, bladder cancer, and breast cancer (Roman-Gomez et al. 2005; Lamprecht et al. 2010; Weber et al. 2010; Wolff et al. 2010; Steidl et al. 2012; Hur et al. 2014; Lock et al. 2014; Miglio et al. 2018).

In 1971, Britten and Davidson proposed that the expansion of transposons in the host genome could have accelerated the evolution of regulatory networks, by acting as a vector to disperse regulatory elements and thereby recruiting host genes into co-regulated, co-expressed networks (Britten and Davidson 1971). Based on this theory, it is expected that regulatory functions, as well as features that account for regulatory activities will be conserved within a group of related transposons. Indeed, transposons within the same subfamilies have been demonstrated to exhibit common regulatory activities. For example, *in silico* analysis has revealed the MER130 DNA transposon subfamily to be significantly enriched within active enhancers in the mouse dorsal cerebral wall (Notwell et al. 2015). These MER130 transposons contain a highly conserved region harbouring putative binding motifs for TFs associated with brain development, and exhibit *in vitro* enhancer activity in mouse embryonic neurons (Notwell et al. 2015). In human innate immunity, LTR retrotransposons are overrepresented in the transcriptional regulatory network (Chuong et al. 2016). In particular, upon interferon-gamma stimulation, the MER41 subfamily is significantly enriched in the binding sites of IRF1 and STAT1, both of which play a crucial role in immune signalling pathways (Chuong et al. 2016). Several MER41 transposons also contribute enhancer activity for immunity-associated genes, such as *AIM2* and *APOL1* (Chuong et al. 2016). In the case of pregnancy, the MER20 subfamily has been found to harbour binding sites for hormone-responsive and pregnancy-related TFs, and are overrepresented near progesterone or cAMP-responsive endometrial genes (Lynch et al. 2011). Epigenetic profiling indicates that these MER20s can exert a diverse range of regulatory activities by acting as enhancers, insulators or repressors (Lynch et al. 2011). As demonstrated by examples above, transposons can exert regulatory activity at the subfamily level, and this conservation of regulatory activity allows functionally correlated genes to be regulated in a temporally or spatially specific manner.

LINE1 retrotransposons are abundant in the human genome, with over 500,000 copies occupying 17% of the genomic sequence (Lander et al. 2001). A full-length LINE1 element is approximately 6kb in length, and contains a 5’ untranslated region (5’UTR), two open reading frames (ORF) and a 3’ poly(A) tail (Figure 1A) (Lander et al. 2001). The 5’UTR of LINE1 transposons harbours a bi-directional promoter (Figure 1A) (Swergold 1990; Speek 2001). LINE1 transposons are classified into subfamilies by the presence of ancestral or post-insertion mutations in their sequences (Boissinot and Furano 2001). L1PA2 transposons are a primate-specific subfamily of LINE1 elements, with approximately 4,940 copies in the human genome (Ovchinnikov et al. 2002). According to RepeatMasker annotations (Karolchik et al. 2004; Smit 2013-2015), 978 out of 4,940 (19.8%) human L1PA2 elements are over 6kb in length, exhibiting limited divergence, deletion and insertion relative to the consensus sequence (Figure 1B).

**Figure 1.**
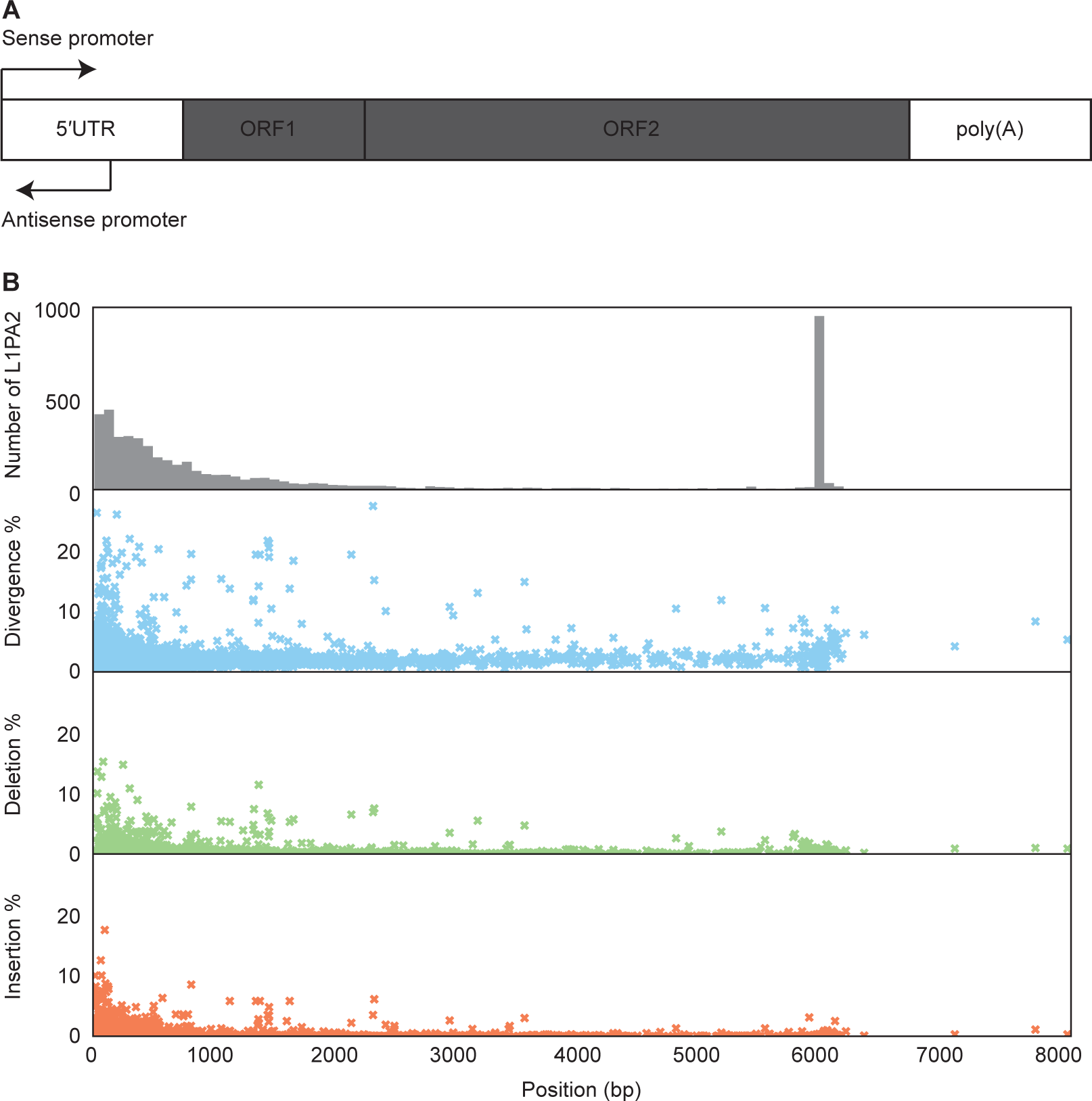
LINE1 structure and human L1PA2 structural summary. **A)** A typical LINE1 transposon, approximately 6kb in length, contains a 5’UTR, two ORFs and a 3’ poly(A) tail. **B)** According to RepeatMasker annotations (Karolchik et al. 2004; Smit 2013-2015), there are approximately 4,940 copies of L1PA2 transposons in the human genome. Top panel displays the length distribution of L1PA2 transposons. The bottom three panels display the percentages of divergence, deletion and insertion against the length of each human L1PA2. Length, degree of divergence, deletion and insertion were calculated using the “genoStart”, “genoEnd”, “milliDev”, “milliDel” and “milliIns” columns from the UCSC RepeatMasker table (Karolchik et al. 2004; Smit 2013-2015).

We previously reported the regulatory activity of several transposon subfamilies in the context of breast cancer (Jiang and Upton 2019). Our analysis revealed that the L1PA2 transposons were significantly enriched in E2F1 and MYC binding sites in MCF7 cells. Furthermore, luciferase assays in triple negative breast cancer cells confirmed the activity of an L1PA2-derived promoter, where the transposon was found to account for a significant proportion of promoter activity to the *SYT1* gene (Jiang and Upton 2019). In support of our findings, a recent paper by Jang *et al*. showed that the L1PA2-derived promoter activity was not limited to *SYT1*, and was also seen for other oncogenes, such as *MET, XCL1* and *AKAP13*, in a number of cancer types (Roman-Gomez et al. 2005; Weber et al. 2010; Hur et al. 2014; Miglio et al. 2018; Jang et al. 2019). In particular, the L1PA2*-MET* alternative transcript has been found in chronic myeloid leukemia, colorectal cancer and breast cancer, and is correlated with enhanced metastasis and poor prognosis (Roman-Gomez et al. 2005; Weber et al. 2010; Hur et al. 2014; Miglio et al. 2018). Intrigued by our results and the literature evidence, we aimed to conduct an integrated investigation on the regulatory activity of the L1PA2 subfamily in breast cancer cells.

Using more comprehensive TF binding data, we showed that the 5’UTR of L1PA2 transposons was a prominent source of TFBSs. In the context of breast cancer, over 27% of L1PA2 transposons contained at least one TFBS in MCF7 breast cancer cells, and the binding of TFs was correlated with active epigenetic modifications. Upon further investigation, several TFs were found to show co-localised binding sites in L1PA2, supporting our hypothesis that L1PA2 could serve as a vector for dispersing functionally related regulatory sequences. *In silico* analysis revealed that the L1PA2-binding TFs formed highly interacting networks, which were functionally enriched for transcriptional mis-regulation in a variety of cancer types. L1PA2 transposons also constituted an abundant reservoir of transcription start sites (TSS) in MCF7 cells. These L1PA2 transposons displayed an active epigenetic profile in MCF7 cells, and contributed cancer-specific promoter activity to a number of alternative or novel transcripts. Taken together, the ubiquitous and replicative nature of the L1PA2 subfamily makes them an exemplary vector for dispersal of co-localised transcription factor binding sites, thereby facilitating the co-ordinated regulation of genes. We demonstrate that the L1PA2 subfamily is a prominent contributor of regulatory elements in breast cancer cells, and likely play a global role in breast cancer transcriptional regulation. The transcriptional activation of L1PA2 transposons in cancer is correlated with alterations in the transcriptome, and may provide novel biomarkers for disease diagnosis and treatment.

## RESULTS

### The L1PA2 5′UTR is a rich reservoir of human TFBSs

To investigate the pattern of TF binding in L1PA2 elements, the occurrence of TFBSs within L1PA2 transposons and neighbouring regions was counted. 2,679 out of 4,940 (∼54.2%) L1PA2 transposons harboured at least one TFBS when all experimental conditions (tissues, cell lines, treatments etc.) were considered. The 5’UTR of L1PA2 transposons harboured abundant TFBSs, as the frequency of TF binding almost doubled in the 5’UTR compared to neighbouring genomic regions (Figure 2A). In contrast, the middle of the transposon sequence had a notable depletion (Figure 2A). This region contains the coding sequences for the LINE1 machinery, and this depletion is consistent with depletion of TF binding within exons (Chen et al. 2015). Decreased mappability within these regions may also have some effects on the appearance of depletion compared to surrounding genomic regions (Sexton and Han 2019). An example of TF binding, as visualised on the GTRD browser (Yevshin et al. 2019), is shown for the L1PA2 transposon located upstream of *MET*, previously described by Roman-Gomez et al. (2005) (Figure 2B).

**Figure 2.**
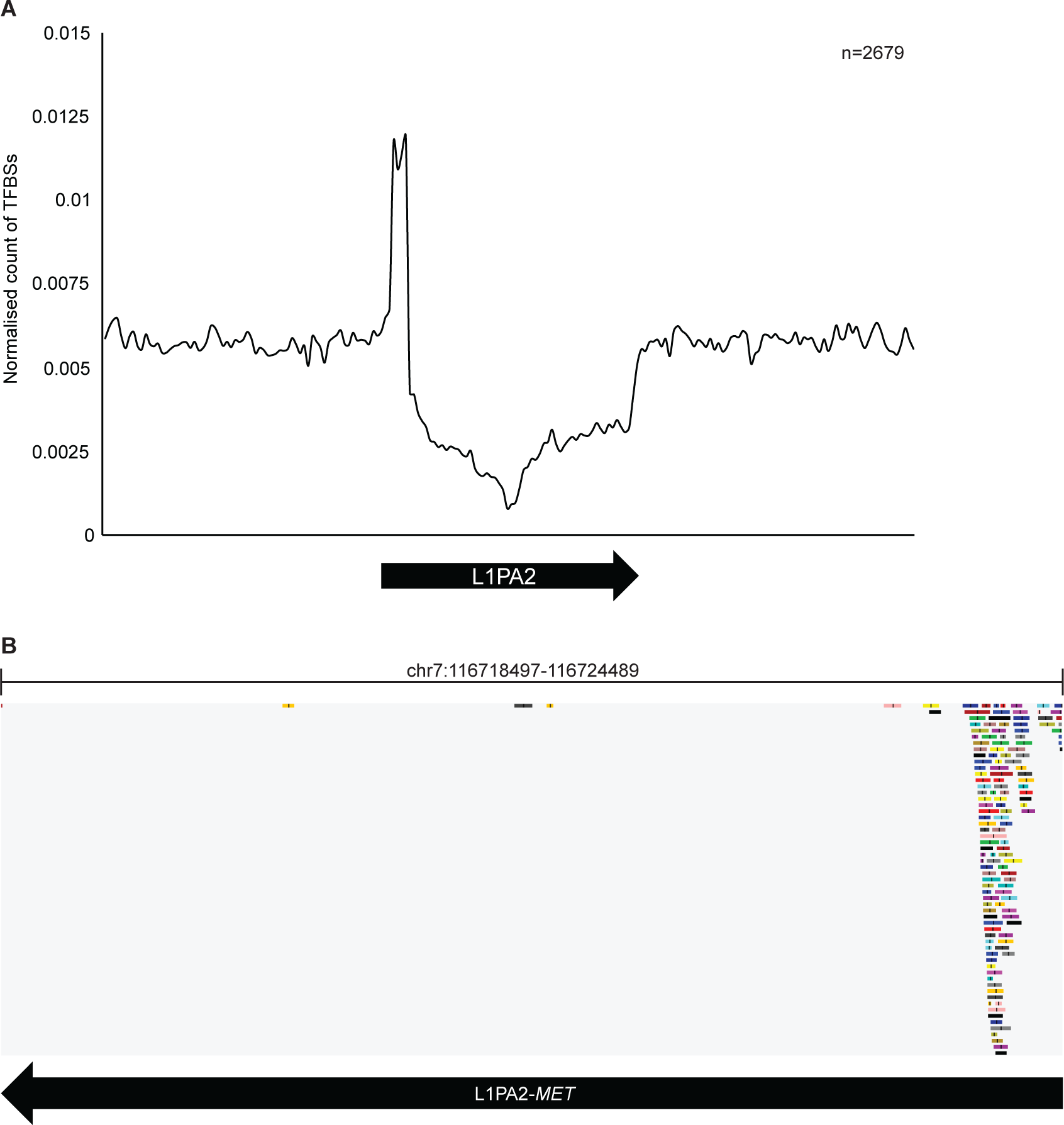
Distribution of TFBSs in human L1PA2 transposons and surrounding regions. **A)** The 5’UTR of L1PA2 was a prominent source of TFBSs. The normalised counts of TFBSs in the 20kb region (bin = 100bp) centred on L1PA2 transposons are shown, averaged by the total number of L1PA2 harbouring at least one GTRD meta cluster (n = 2,679). While the majority of L1PA2 structures showed a depletion of TFBSs, the 5’UTR was a prominent reservoir of TFBSs. **B)** Example GTRD browser view of TFBS distribution in L1PA2 transposons. The TF binding distribution in the L1PA2 transposon located upstream of the *MET* gene was visualised on the GTRD browser (Yevshin et al. 2019). Each rectangular track indicates a TF binding meta-cluster. Black arrows indicate the position and orientation (5’ to 3’) of the L1PA2 transposons.

### Over 27% of L1PA2 transposons harbour TFBSs in a breast cancer cell line

Next we analysed cell-type-specific ChIP-seq data to investigate the TF binding status of L1PA2 transposons in the context of breast cancer. A total of 1,376 out of 4,940 (27.9%) L1PA2 elements harboured at least one TFBS in MCF7 breast cancer cells alone.

To investigate whether this binding was the result of functional or spurious binding events, we investigated the DNAse sensitivity of L1PA2 transposons via the analysis of DNA-seq datasets. DNAse hypersensitivity assays provide a proxy for open chromatin regions, but do not necessarily indicate active transcription (Meyer and Liu 2014; Tsompana and Buck 2014). There is a general positive correlation between DNAse signal and transcriptional activity (Zhang et al. 2012). Interestingly, DNAse hypersensitivity was observed in the 5’UTR of both bound and unbound elements; however, the strength of this signal was around twice as strong for bound elements, supporting a more open chromatin configuration and transcriptional activity from bound transposons (Figure 3A).

**Figure 3.**
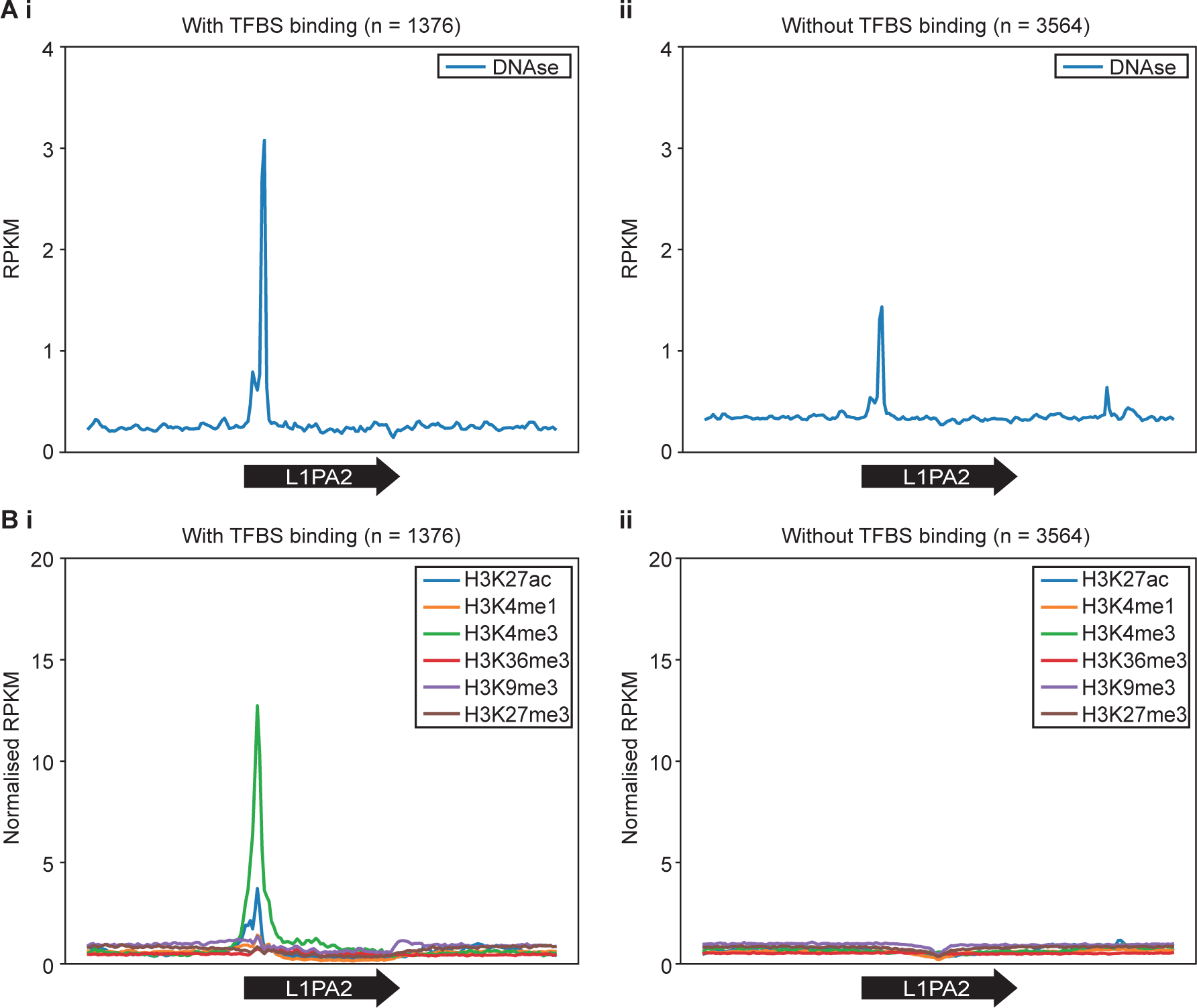
TF binding in L1PA2 was correlated with **A)** DNAse sensitivity and **B)** active histone tail modifications. The RPKM values of the DNAse-seq data and the average RPKM values of histone modification ChIP-seq data are shown for the 20kb region (bin = 100bp) centred on L1PA2 transposons **i)** with, and **ii)** without TFBSs. Black arrows indicate the position and orientation (5’ to 3’) of L1PA2 transposons.

To further confirm this apparent epigenetic activation and TF binding, we investigated the histone states of L1PA2 elements in MCF7 cells by analysing publicly available ChIP-seq datasets. We reasoned that functional binding would be associated with active chromatin marks commonly observed in promoters and enhancers (H3K27ac, H3K4me1, and H3K4me3), or gene bodies (H3K36me3), but not repressive chromatin marks (H3K9me3 and H3K27me3) (Peters et al. 2003; Barski et al. 2007; Creyghton et al. 2010).

Unbound L1PA2 elements were not remarkably different from their surrounding genomic regions, suggesting these elements were transcriptionally inactive as expected, and provided a baseline expectation (Figure 3B). In contrast, we found that active histone modifications were enriched in the 5’ UTR of TF-bound L1PA2 elements, indicating these elements were transcriptionally active (H3K27ac), with a strong promoter profile (H3K4me3), and some elements potentially contained primed enhancer properties (H3K4me1) (Figure 3B). No notable features were observed for H3K36me3 histone modifications that usually mark active gene bodies (Figure 3B). A slight increase in the repressive H3K9me3 marks was seen in the regions flanking the transposon 5’ ends, which may suggest there was a counteracting mechanism attempting to repress ectopic transposon activation (Figure 3B). Taken together, this data strongly supports the existence of L1PA2-derived promoters and potentially L1PA2-derived enhancer elements in MCF7 breast cancer cells.

### L1PA2 contains co-localising TFBSs in MCF7 cells

A total of 75 TFs, including ESR1, SFPQ and MYC, were found to bind to L1PA2 transposons in MCF7 cells (Figure 4A). We sought to investigate whether these TFs were known to bind in a co-ordinated manner, which is an essential feature of transcriptional complexes (Wasserman and Sandelin 2004; Harbeck et al. 2019). We confirmed that in MCF7 cells, a number of L1PA2 transposons showed a consistent binding pattern where they were found to harbour co-localised TF binding sites (Figure 4B). More specifically, binding sites of TFs with the highest binding frequency (ESR1, SFPQ, MYC, FOXA1, NR2F2, CTCF, E2F1, KDM5B and ZNF143) predominantly mapped to the 5’UTR of L1PA2s, and these binding sites often occurred within close proximity to each other (Figure 5).

**Figure 4.**
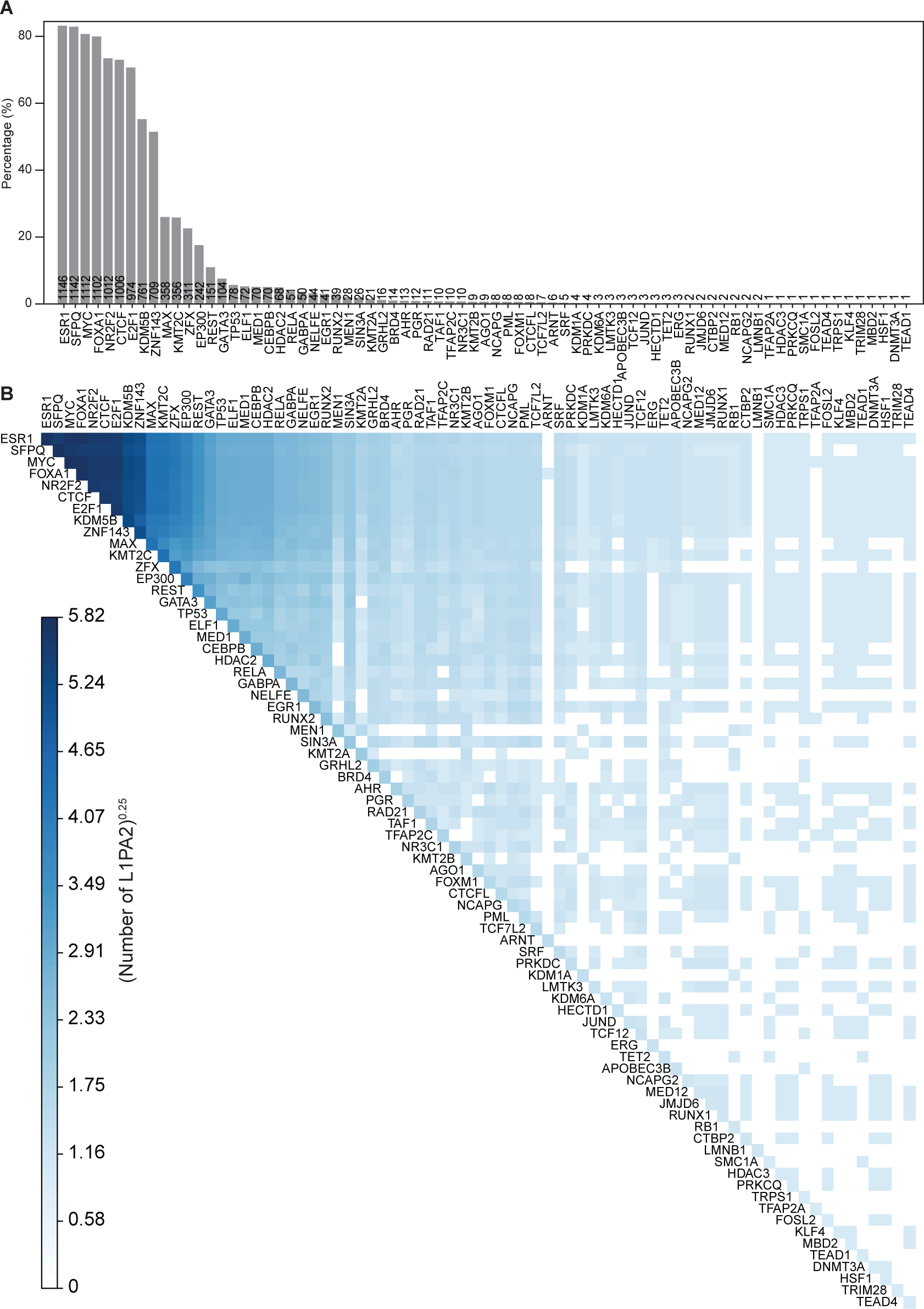
L1PA2 contained co-localising TFBSs in MCF7 cells. **A)** A total of 75 TFs were found to bind to L1PA2 transposons in MCF7 cells. The bars and numbers show the percentages and counts of TF-bound L1PA2 transposons (n = 1,376) respectively. **B)** For each pair of TFs, the number of L1PA2 transposons containing both TFBSs is shown in the heatmap, where the counts are transformed (n^0.25^).

**Figure 5.**
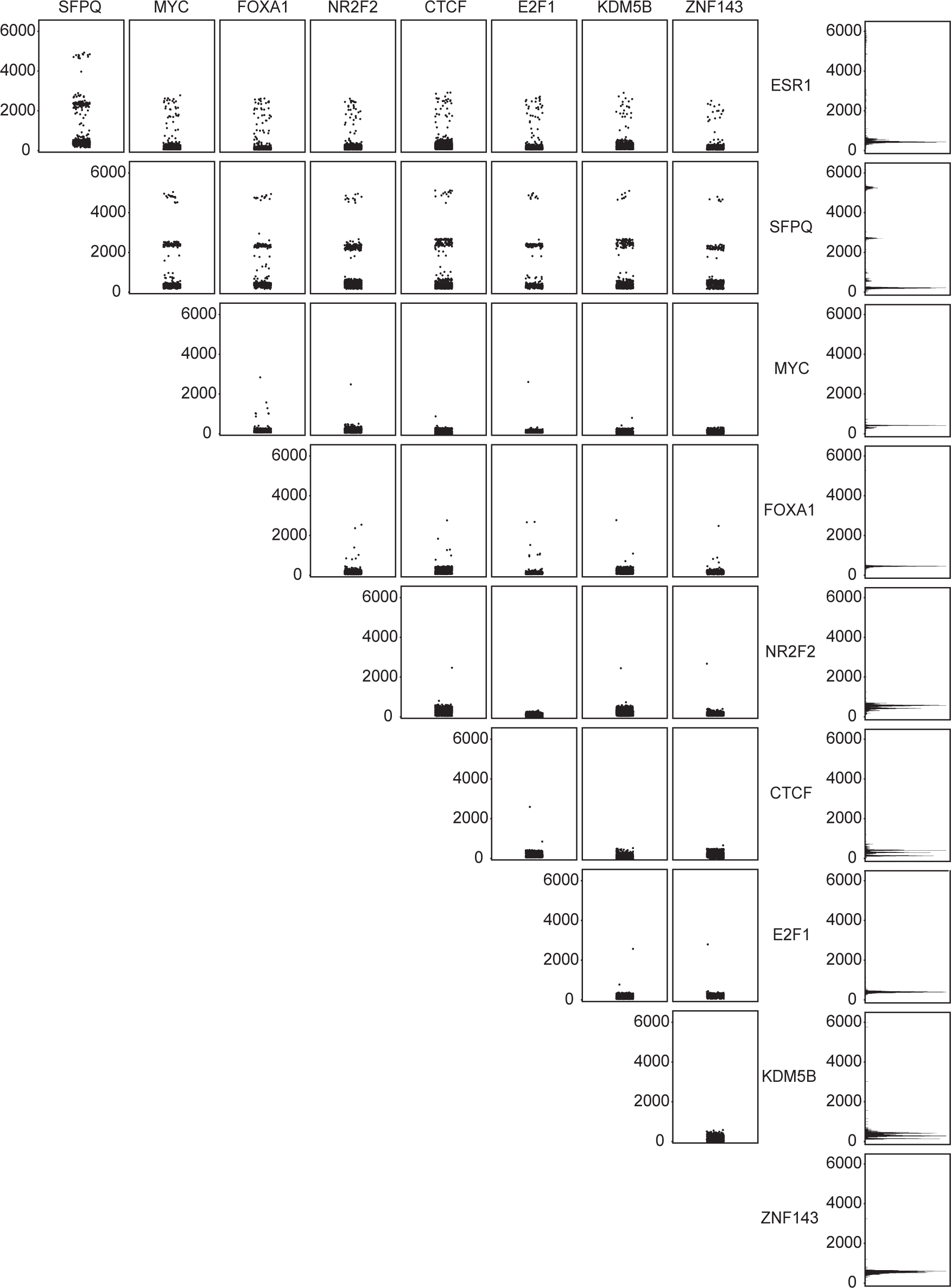
The majority of TFBSs co-localised to the 5’UTR of L1PA2s. Pairwise strip plots (left) indicate the distances (bp) between the binding sites of each pair of TFs found within the same L1PA2 transposons, and the bar graphs (right) indicate the position (bp) of TFBSs for each TF in the consensus L1PA2 transposon.

### TFBS co-localisation occurs more often in L1PA2 compared to the rest of the genome

We next compared the occurrence of TF binding site co-localisation in L1PA2 transposons and genome-wide. Asymmetrical co-localisation was common when considering genome-wide TFBSs, possibly due to the variations in the total numbers of binding sites (Figure 6A). For example, while nearly all NR2F2 binding sites (93%) co-localised with ESR1 binding sites, only 23.2% of ESR1 sites co-localised with NE2F2 sites. This asymmetry can be seen as a function differences in the total number of binding sites for each factor, where ESR1 binding sites were four times more abundant than NE2F2 binding sites.

**Figure 6.**
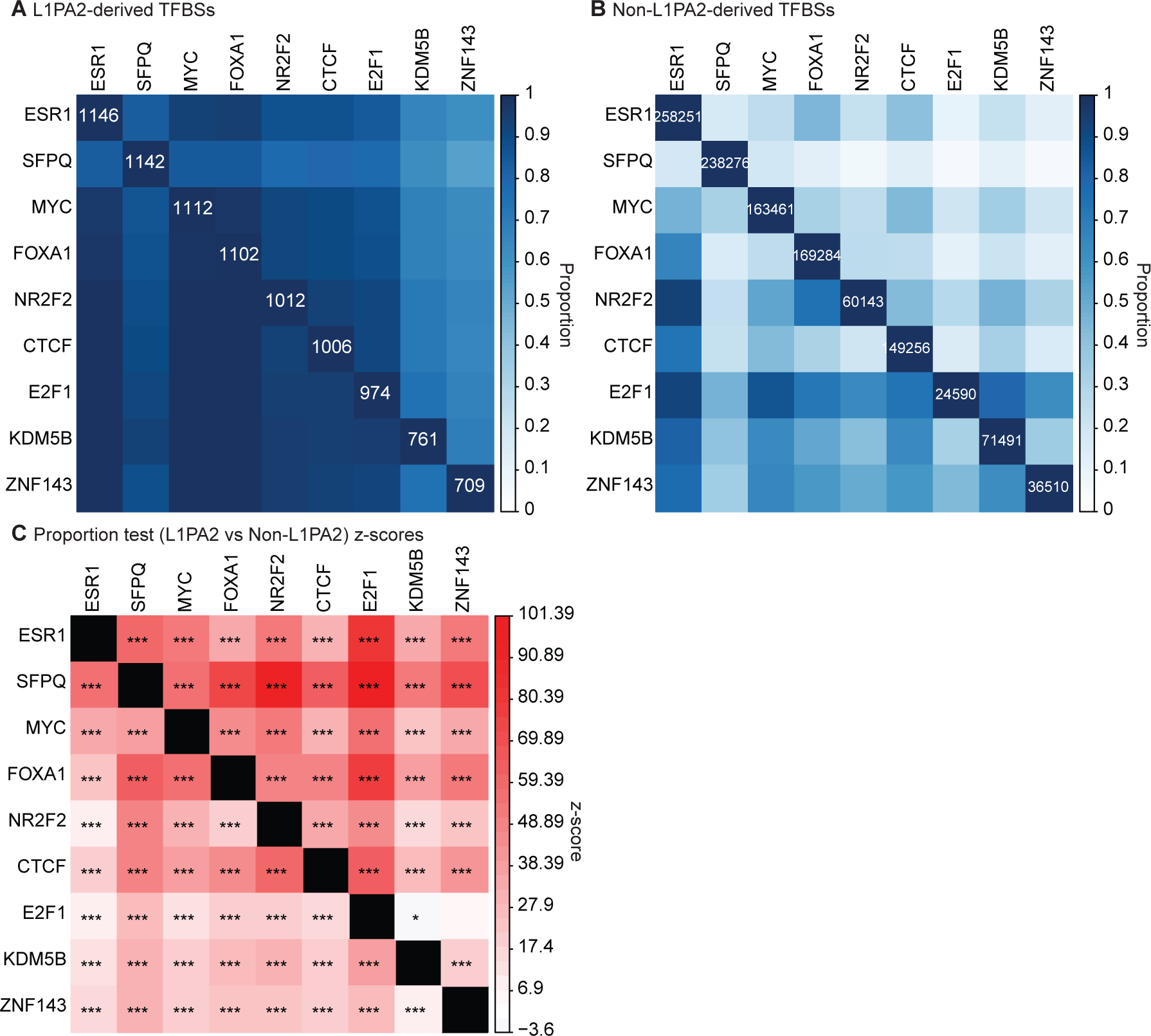
Top binding TFs co-occurred more frequently in L1PA2s than in the rest of the genome. The proportions of co-occurrence are shown for **A)** L1PA2-derived and **B)** non-L1PA2-derived TFBSs. For each TF, the total number of binding sites is shown in the diagonal. **C)** For each TF-TF combination, the co-localisation proportions were compared (L1PA2 versus non-L1PA2) using a one-tailed proportion z-test. The heatmap indicates the z-scores of the proportion tests, where a positive z-score (red) indicates more co-localisation in L1PA2s, and a negative z-score (blue) indicates more co-localisation in the rest of the genome. P < 6.2E-4: *, p < 0.0001: **, p < 0.00001: ***. To account for asymmetrical co-localisation, the proportions and z-scores correspond to the chances of the TF in the row co-localising with the TF in the column.

In contrast, the total numbers of L1PA2-derived TFBSs were more similar amongst the top binding TFs, which led to less asymmetrical co-localisation, and supported the hypothesis that these TFBSs co-occurred more in L1PA2 transposons (Figure 6B). Indeed, proportion tests revealed that, overall, the binding sites of top binding TFs co-occurred more frequently in L1PA2 transposons when compared to the rest of the genome (Figure 6C). The only exceptions involve E2F1, where it appeared to co-localise with KDM5B less frequently in L1PA2, and displayed no significant enrichment or depletion for ZNF143 co-localisation.

### L1PA2 contains binding sites for highly interacting, cancer-associated TF networks in MCF7 cells

We interrogated known biological observations of protein-protein interactions, and performed functional enrichment analysis to identify biological functions and disease states associated with the L1PA2-binding TFs.

STRING analysis showed that these 75 TFs formed highly interacting networks, based on available experimental evidence (PPI enrichment p-value < 1E-16) (Szklarczyk et al. 2019) (Figure 7A). Functional enrichment analysis by ToppFun revealed that these TFs were significantly enriched in 44 pathways, with the most significant enrichment reported in transcriptional mis-regulation in cancer (Bonferroni q-value = 7.08E-10) (Chen et al. 2009) (Figure 7B). For the complete list of significant pathways see Supplementary Table 1.

**Figure 7.**
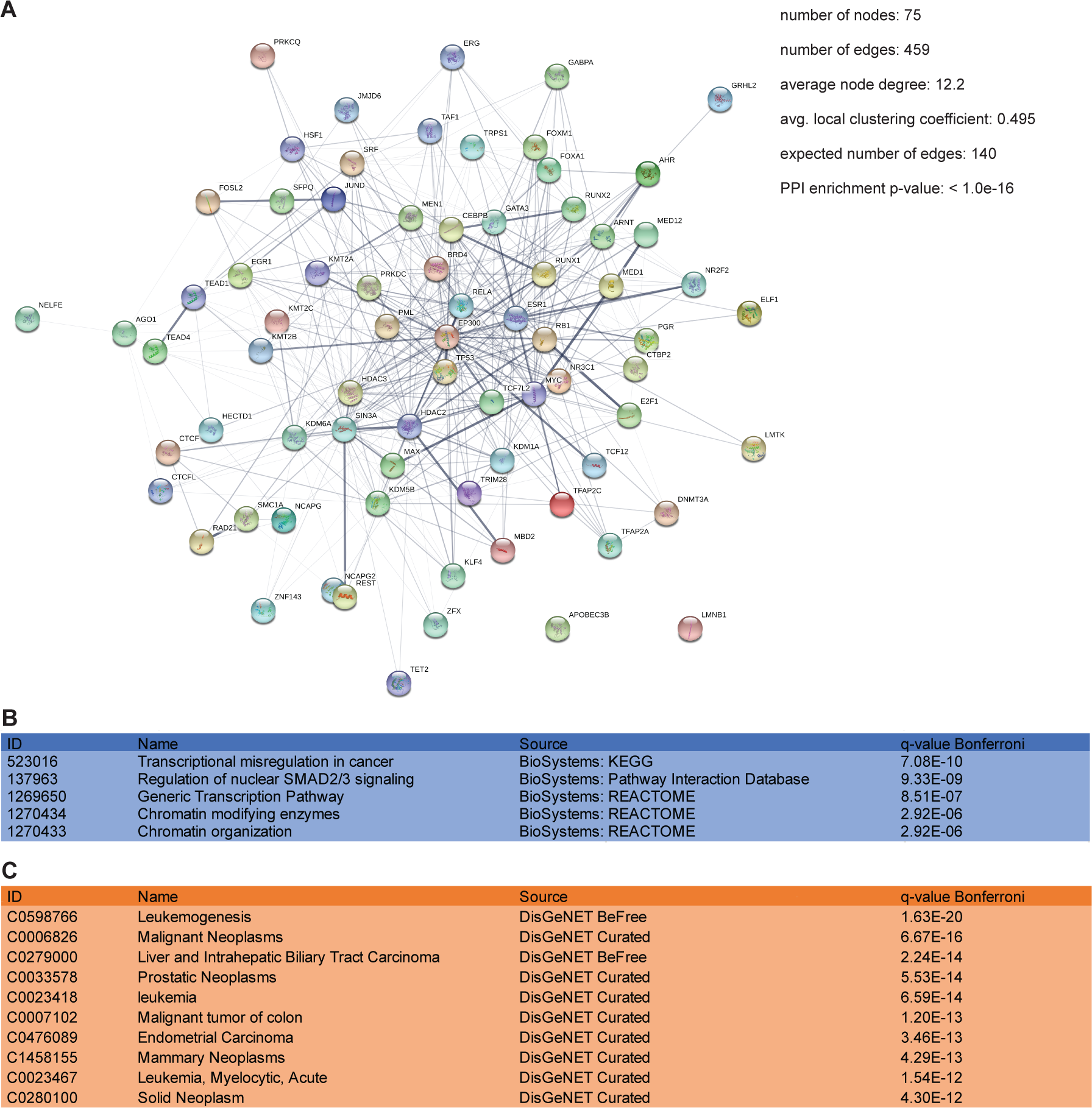
L1PA2 contained binding sites for highly interacting, cancer-associated TF networks in MCF7 cells. **A)** L1PA2-binding TFs were predicted to form highly interacting networks according to STRING analysis (Szklarczyk et al. 2019). Edges represent protein-protein interactions with experimental evidence, and the thickness of edges represents level of confidence. Statistical analysis results from STRING are shown. **B)** Pathway enrichment analysis by ToppFun showed that L1PA2-binding TFs were enriched for transcriptional mis-regulation in cancer (Chen et al. 2009). Top enriched pathways are shown. **C)** Disease enrichment analysis by ToppFun showed that L1PA2-binding TFs were enriched in various cancer types (Chen et al. 2009). Top enriched diseases are shown.

Considering diseases, ToppFun analysis showed that these TFs were significantly enriched in 228 diseases, 183 of which are neoplastic diseases, including prostate cancer (prostatic neoplasms, ranked 4^th^) (Bonferroni q-value = 5.53E-14) and breast cancer (mammary neoplasms, ranked 8^th^) (Bonferroni q-value = 4.29E-13) (Chen et al. 2009) (Figure 7C). For the complete list of significant diseases see Supplementary Table 2.

### Oncogenic TF binding motifs are conserved in the L1PA2 5’UTR

We hypothesised that the transcription factor binding activity shared amongst L1PA2 transposons was correlated with conserved motif sequences, and performed motif analysis to identify the binding motifs of frequently binding, oncogenic TFs (ESR1, FOXA1 and E2F1) in L1PA2 sequences. Overall, 66.8%-98.3% of TF-bound L1PA2 transposons contained at least one significant motif of the corresponding TF (Figure 8). The majority of binding motifs mapped to the 5’UTR of L1PA2 elements (Figure 8). In particular, four ESR1 binding motif peaks were identified in the 5’UTR, and each peak contained a highly consistent, overrepresented sequence, which was similar to the palindromic estrogen response element. For FOXA1, the majority of the discovered motifs were located in the 5’UTR, with a small number mapping to the 3’ end of L1PA2 transposons. E2F1 binding motifs showed the highest level of conservation, with highly consistent motifs identified almost exclusively in the 5’UTR.

**Figure 8.**
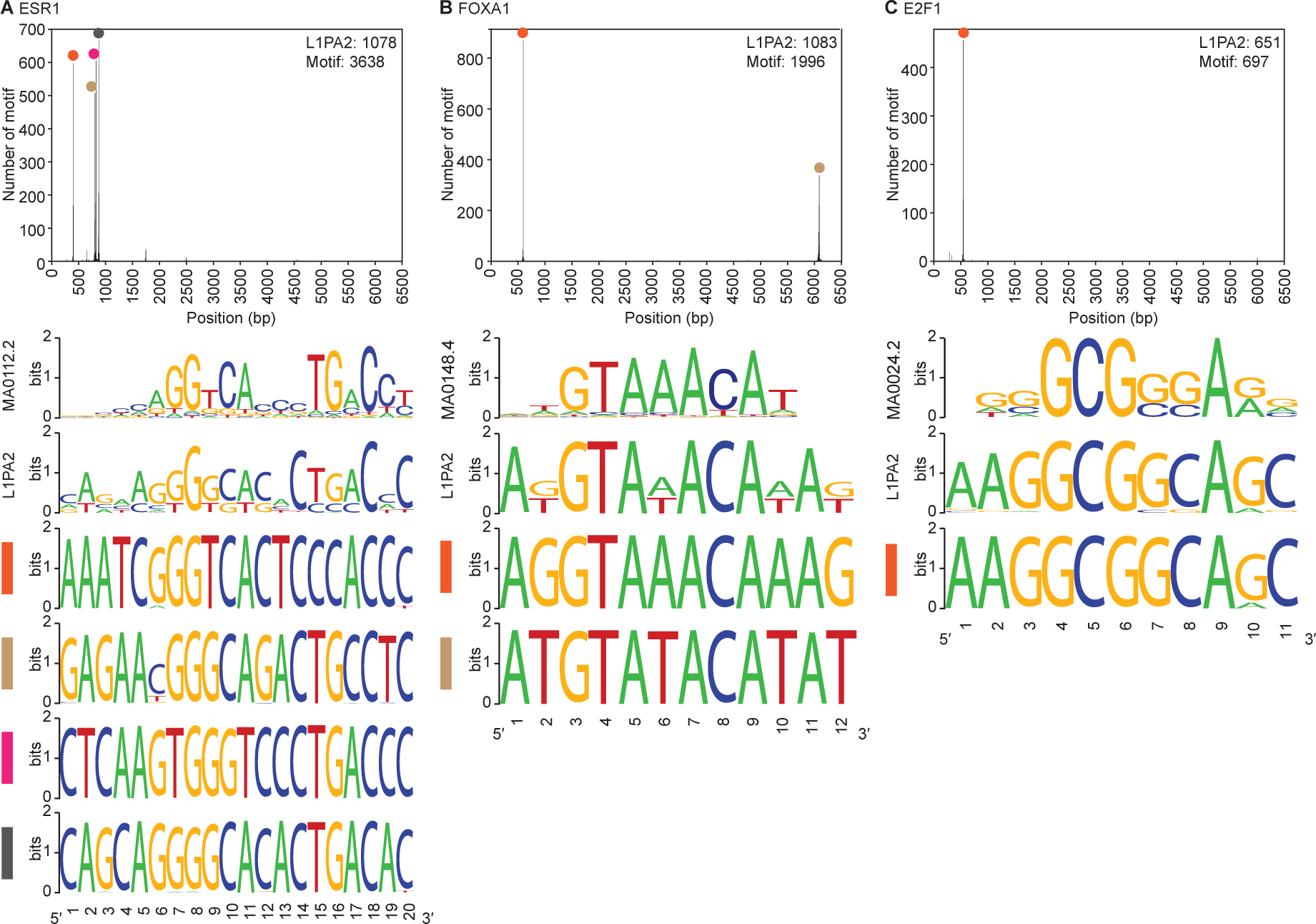
Oncogenic TF binding motifs are conserved in the 5’UTR of L1PA2 transposons. Motif analysis was performed to identify the binding motifs for **A)** ESR1, **B)** FOXA1 and **C)** E2F1 in L1PA2 transposons. The positions of the discovered motifs in the consensus L1PA2 sequence are shown in the histograms (bin = 5bp). The number of L1PA2 elements containing at least one statistically significant motif, and the total number of statistically significant motifs are shown. For each TF, the JASPAR sequence logo is shown, labelled with the corresponding JASPAR motif ID. The “L1PA2” sequence logo was generated using all discovered motif sequences. Each peak in the histogram is labelled with a colour, and the sequence logos generated from motifs in the corresponding bins are shown.

### TF binding in L1PA2 transposons is correlated with the activation of nearby genes

The promoter regions of genes, which harbour crucial regulatory sequences, are usually found within 1kb of the TSSs, while other regulatory elements, such as enhancers, are typically located farther away from their target genes (Vernimmen and Bickmore 2015). To investigate the regulatory role of L1PA2-derived TF binding, the distribution of L1PA2 transposons with respect to breast cancer-associated genes was analysed using publicly available RNA-seq data from MCF7 breast cancer and MCF10A near-normal cells. 9,387 and 9,781 transcripts were up-regulated and down-regulated in MCF7 cells respectively. Overall, TF-bound L1PA2 transposons were correlated with the activation of breast cancer genes in MCF7 cells, as a higher percentage of TF-bound L1PA2 transposons were found in the promoter regions, as well as up to 20kb away from the TSSs of up-regulated transcripts (Figure 9). In contrast, TF-bound L1PA2 transposons were found less frequently in the promoter or surrounding 20kb regions of down-regulated transcripts (Figure 9). Similarly, L1PA2s that lacked TF binding were generally absent from the promoter region of differentially expressed transcripts (Figure 9).

**Figure 9.**
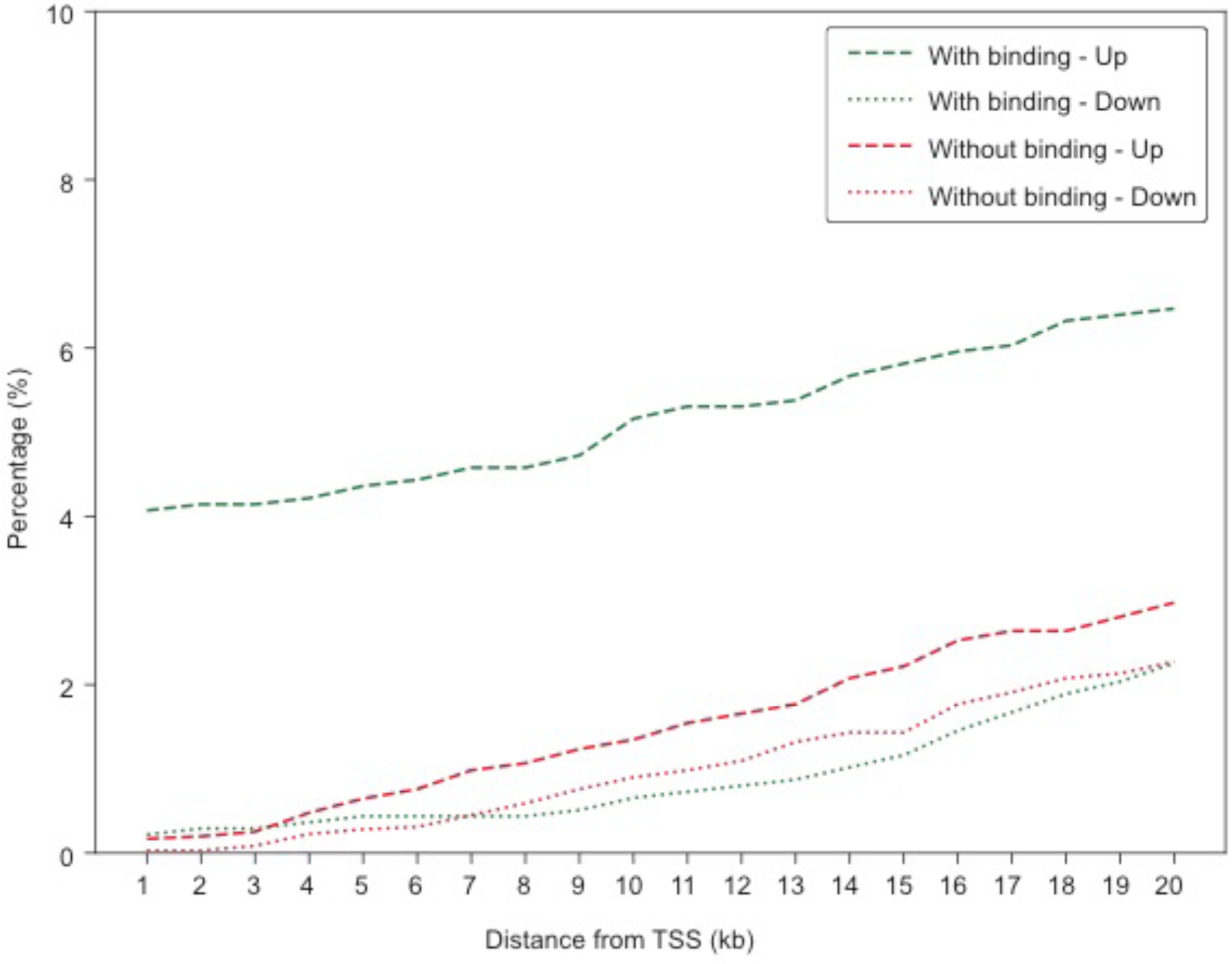
TF binding in L1PA2 transposons was correlated with activation of neighbouring genes. The percentages of transposons located up to 20kb away from the TSSs of differentially expressed transcripts are shown for L1PA2 transposons with (green) and without (red) binding. Dash and dotted lines indicated up-regulated and down-regulated transcripts respectively.

### L1PA2 transposons are overrepresented in the TSSs of up-regulated genes

To assess the contribution of L1PA2 transposons to transcript initiation, direct overlap between L1PA2 and the TSSs of differentially expressed transcripts was analysed. 41 out of 1,376 TF-bound L1PA2 transposons harboured the TSSs of up-regulated transcripts, while only one L1PA2 harboured the TSS of a down-regulated transcript. Enrichment analysis by random rotation of the genome showed that L1PA2 transposons were significantly enriched for the TSSs of up-regulated transcripts (p = 4.83E-11), and significantly depleted for down-regulated transcripts (p = 2.69E-09). L1PA2 transposons harbouring TSSs of up-regulated transcripts are summarised in Supplementary Table 3. For results of transcript expression quantification by StringTie (Pertea et al. 2015) see Supplementary Figure S1.

### L1PA2 transposons bearing up-regulated TSSs exhibit active epigenetic marks in MCF7 cells

We further confirmed the cancer-specific transcriptional activity of TSS-bearing L1PA2 transposons by comparing their epigenetic states in MCF7 cells and MCF10A cells. Epigenetic profiling revealed that TSSs in L1PA2 transposons were correlated with increased active histone tail modifications and increased DNAse sensitivity in MCF7 cells, while these active marks were not observed for the same regions in MCF10A cells (Figure 10A). These L1PA2 transposons also contained RNA pol II binding sites previously reported in breast cancer cell lines (Figure 10A). The exon-intron structure of the L1PA2-derived transcripts, as well as the corresponding known transcripts, are shown in Figure 10B.

**Figure 10.**
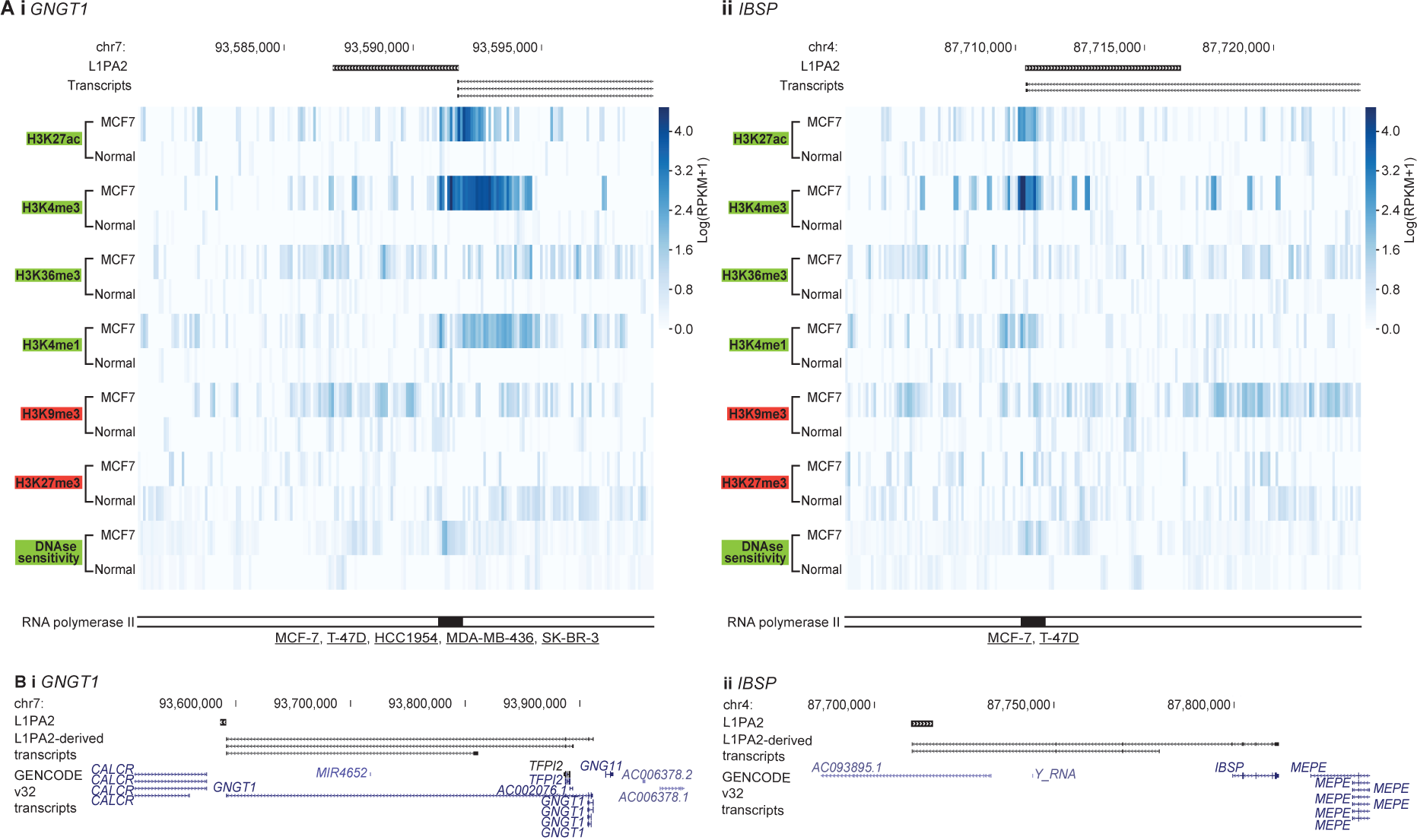
L1PA2 bearing up-regulated transcript TSSs showed cancer-specific, active epigenetic profiles. Examples are shown for **i)** *GNGT1* and **ii)** *IBSP*. For each gene, **A)** histone tail modifications and DNAse sensitivity in MCF7 and normal tissues are shown for the 20kb region centred on the L1PA2. RNA pol II binding within each L1PA2, as well as the cell lines where the binding was detected, are shown. Underlined cell line names indicate cancerous cell lines. **B)** The exon structures of the L1PA2-derived transcripts (black), compared to the GENCODE v32 transcripts (blue), are shown (Frankish et al. 2019).

## DISCUSSION

Approximately 45% of the human genome is composed of transposons (Lander et al. 2001). For transposons to integrate and mobilise within a host genome, by necessity they must contain cis-regulatory sequences that are compatible with host RNA polymerases and/or TFs (Jordan et al. 2003; Cruickshanks and Tufarelli 2009; Gifford et al. 2013; Sundaram et al. 2014; Chuong et al. 2016; Chuong et al. 2017). This compatibility also enables the host to exapt the transposon-derived regulatory sequences to modify the transcriptional regulation of host genes, including the generation of novel transcript isoforms originating in transposon-derived promoters.

Transposons have been exapted to contribute regulatory roles in diverse biological processes, including innate immunity and pregnancy (Lynch et al. 2011; Sundaram et al. 2014; Chuong et al. 2016). In some cases, multiple elements of a given transposon subfamily have been exapted to contribute to the co-ordinated regulation of multiple genes, such as the MER41 subfamily that contributes enhancer activities to immunity-related genes such as *AIM2, APOL1, IFI6*, and *SECTM1* (Chuong et al. 2016). By replicating highly related sequences throughout the genome, transposons provide abundant substrate for exaptation to mediate the temporal or spatial co-ordinated regulation of functionally related genes.

Individual L1PA2 transposons have been reported to modulate gene expression in various cancer types. More specifically, a hypo-methylated L1PA2 transposon is found to act as an alternate promoter to the *MET* oncogene in breast cancer, chronic myeloid leukemia and colorectal cancer, and the L1PA2*-MET* expression is associated with enhanced malignancy and poor prognosis (Roman-Gomez et al. 2005; Weber et al. 2010; Hur et al. 2014; Miglio et al. 2018). In the context of breast cancer, we previously identified enriched E2F1 and MYC binding sites in the L1PA2 subfamily, and demonstrated that L1PA2 contributed the significant promoter activity for the *SYT1* oncogene in triple negative breast cancer cell lines (Jiang and Upton 2019). Intrigued by these results, we hypothesised that this regulatory activity was conserved across the L1PA2 subfamily, and performed an integrated *in silico* analysis of L1PA2-derived TF binding and promoter activity in breast cancer cells.

Here we have utilised data from the GTRD and the ChIP-Atlas databases, both of which aim to catalogue and compile an extensive range of publicly available ChIP-seq datasets (Oki et al. 2018; Yevshin et al. 2019). The GTRD data was used for general analysis of TF binding, as it merges the binding sites for individual TFs into meta-clusters representing all experimental conditions (tissues, cell lines, treatments etc.), reducing the computational resources required for analysis. By contrast, the ChIP-Atlas database retains the cell line and tissue information, allowing us to assess TF binding in breast cancer cell lines specifically.

Our analysis revealed that when all TFs and experimental conditions in the GTRD database were considered, 54.2% of L1PA2 transposons contained at least one TFBS, and TFBSs often clustered within the 5’UTR (Figure 2). This represents a minimal estimation of L1PA2 regulatory potential, as many TFs have been studied under a limited number of conditions. The 5’UTR of LINE1 transposons, approximately 900bp in length, contains two internal promoters allowing transcription to be initiated in either direction (Swergold 1990; Speek 2001). This region represents a potent reservoir of regulatory activity, harbouring binding sites for a variety of TFs, including YY1, RUNX, SOX and MYC (Becker et al. 1993; Tchénio et al. 2000; Yang et al. 2003; Sun et al. 2018). The LINE1 5’UTR, particularly the antisense promoter, also contributes significant promoter activity in a variety of cell types, affecting the expression of many genes such as *SYT1, MET* and *JAK1* (Criscione et al. 2016; Jiang and Upton 2019).

To focus our analysis on the regulatory activity of L1PA2 in breast cancer, we investigated the TF binding activity of L1PA2 in MCF7 breast cancer cells. Approximately 27.9% of L1PA2 transposons were bound by at least one TF in MCF7 cells, and the binding was correlated with active epigenetic marks in the 5’UTR, suggesting potential promoter or enhancer activities of these transposons (Figure 3). A total of 75 TFs bound to L1PA2 transposons, including known oncogenic TFs such as ESR1, FOXA1 and E2F1 (Figure 4A) (Zacharatos et al. 2004; Horiuchi et al. 2012; Lei et al. 2019). Motif analysis revealed that the binding of these oncogenic TFs was correlated with conserved sequence motifs in the 5’UTR of L1PA2 transposons (Figure 8).

In particular, the expression of the estrogen receptor (ESR1 or ER), is the defining feature of the Luminal A and Luminal B breast cancer subtypes (Harbeck et al. 2019). At least 70% of breast cancer cases are ER-positive, and ER signalling is a key driver of cancer progression and tumour growth (Harbeck et al. 2019; Lei et al. 2019). Amongst all L1PA2-binding TFs in MCF7 cells, ESR1 shows the highest binding frequency, with over 80% of TF-bound L1PA2s harbouring an ESR1 binding site (Figure 4A). The conserved ESR1 binding in L1PA2 suggests that the L1PA2 subfamily likely plays a prominent role in mediating ESR1 regulation.

In 1971, Britten and Davidson proposed that transposons could act as a vector to disperse regulatory elements and thereby recruit genes into co-regulated, co-expressed networks (Britten and Davidson 1971). Intrigued by their theory, we sought to investigate whether L1PA2 transposons can act as a vector for dispersing regulatory elements that contribute to breast cancer. We analysed the frequency of co-localisation between each pair of top L1PA2-binding TFs in MCF7 cells, and found that the binding sites of these TFs indeed co-occurred significantly more frequently in L1PA2 transposons, particularly in the 5’UTR, compared to the remainder of the genome (Figure 4B, Figure 5&6). In particular, the binding sites of ESR1, which binds DNA upon estrogen stimulation, was found to co-localise with the binding sites of E2F1 and MYC, both of which play a role in transcriptional regulation of estrogen-stimulated genes (Cheng et al. 2006; Stender et al. 2007). Furthermore, MYC and CTCF also co-localised to the 5’UTR of L1PA2 transposons, which was a pattern that has also been observed for the L1HS subfamily, and may be related to chromatin remodelling (Sun et al. 2018). Overall, the enrichment of co-occurring TFBSs in L1PA2 transposons supported the hypothesis that L1PA2 transposons served as a vector for dispersing functionally related regulatory sequences, and thereby mediated the combinatorial control of genes in breast cancer.

Mammalian gene regulation is effected by the co-ordinated action of multiple transcription factors, acting in transcriptional complexes (Wasserman and Sandelin 2004; Harbeck et al. 2019). Therefore, understanding the activity of interacting TF networks, rather than individual TFs, often provides a more comprehensive insight into gene regulation in the cancer transcriptome. Considering only protein-protein interactions with experimental evidence in the STRING database (Szklarczyk et al. 2019), L1PA2-binding TFs formed a highly interacting network, demonstrating that these TFs commonly interact in gene regulation (Figure 7A). Functional enrichment analysis by ToppFun (Chen et al. 2009) supported the oncogenic properties of these networks, as the L1PA2-binding TFs showed a statistical enrichment for transcriptional mis-regulation in multiple cancer types, including prostate cancer and breast cancer (Figure 7B&C). Taken together, L1PA2 transposons harbour binding sites for functionally interacting, cancer-associated TFs, and contribute to co-ordinated transcriptional regulation in breast cancer.

Transcriptional regulation is often a complex and highly dynamic process, and can lead to both the activation or repression of neighbouring genes depending on the interactions of TFs. To better understand the regulatory impact of L1PA2 transposons in breast cancer, we re-analysed publicly available RNA-seq data from MCF7 breast cancer cells and MCF10A near-normal cells, and assessed the distribution of L1PA2 relative to up-regulated (activation) or down-regulated (repression) transcripts in MCF7 cells. TF binding in L1PA2 transposons was correlated with the activation of nearby transcripts (Figure 9). Detailed assessment of TSS and L1PA2 overlap showed that the up-regulated transcripts were significantly enriched in L1PA2 transposons, and that the L1PA2 subfamily was a major driver of transcript expression in MCF7 cells. Some of these L1PA2-derived TSSs have been annotated in the Ensembl database (Aken et al. 2016), while many appeared to be alternative or novel TSSs that were not yet annotated (Supplementary Table 3). Some L1PA2-derived transcripts shared exons with known transcripts, such as *GNGT1* and *IBSP*, representing novel isoforms (Figure 10B). These alternate transcripts were likely products of L1PA2 epigenetic dysregulation in the cancer state, supported by increased active histone tail modifications and increased DNAse-sensitivity (Figure 10A). Further supporting the cancer-specific regulatory activity of these L1PA2s, RNA pol II binding sites within the L1PA2s were found, and these binding sites have previously been reported, sometimes exclusively, in cancer cell lines (Figure 10A). Amongst these L1PA2-derived up-regulated transcripts, we found the L1PA2-*MET* transcript, which had previously been found to be linked to enhanced malignancy and poor prognosis (Roman-Gomez et al. 2005; Weber et al. 2010; Hur et al. 2014; Miglio et al. 2018) (Supplementary Table S3). Taken together, transposons that harbour TFBSs are transcriptionally active, and provide novel TSSs to unannotated transcript isoforms in MCF7 cells. Some of these transcripts have established oncogenic properties, and it is likely that the other L1PA2-driven up-regulated transcripts, summarised in Supplementary Table 3, also have oncogenic functions in breast cancer.

In this study, we performed an integrated analysis to assess the regulatory activity of the L1PA2 transposon subfamily in the MCF7 breast cancer cell line. Our analyses demonstrated that L1PA2 transposons, primarily their 5’UTR, constituted a rich reservoir of regulatory potential, as they contributed abundant functional TFBSs in various tissues and experimental conditions. In the context of breast cancer, L1PA2 transposons facilitated binding site co-localisation of functionally interacting TFs, and thereby contributed to the co-ordinated gene regulation in cancer. The transcriptional activation of L1PA2 transposons in breast cancer cells, potentially due to loosened epigenetic repression, led to alterations in the transcriptome and the production of many alternative or novel transcripts. Some of these L1PA2-driven transcripts, such as L1PA2-*MET*, have well established oncogenic properties, and it is likely that the other transcripts also exhibit similar oncogenic activities in cancer. In summary, L1PA2 transposons contribute abundant regulatory sequences for the co-ordinated gene regulation and oncogene activation in breast cancer. LINE1 transposons, including L1PA2, have been touted as potential cancer biomarkers, but have not been successfully adapted for such a use. By contrast, transcripts resulting from their transcriptional activation may serve as proxies for dysregulated transposons in tumour cells, and provide novel biomarkers for cancer detection. While further investigation is required on the function of additional putative oncogenes, these may also provide targets for therapy or disease prognosis. Furthermore, targeting L1PA2 transcriptional activation may itself provide a broad-acting target to control aberrant gene regulation in breast cancer.

## METHODS

### Genomic locations of genetic entities

Genetic entities that mapped to alternate chromosomes, including TFBSs, transposons, and transcripts, were excluded from all analyses in this study. All genomic locations were in hg38 unless otherwise stated.

### Identifying TFBSs in L1PA2 and surrounding regions

TFBSs, defined as meta-clusters, were acquired from the GTRD database (version 19.10) (Yevshin et al. 2019). The GTRD v19.10 database includes ChIP-seq datasets performed in 609 human cell lines and 289 tissue types, and merges binding sites identified across various experimental conditions (tissues, cell lines, treatments etc.) into meta-clusters (Yevshin et al. 2019). The genomic locations (hg38) of human L1PA2 elements were retrieved from the UCSC RepeatMasker table (Karolchik et al. 2004; Smit 2013-2015). For each L1PA2 element, a 20kb region centred on the transposon was defined and divided into 100bp bins. The genomic locations of the GTRD TFBSs were intersected with the L1PA2 elements using BedTools Intersect, where an overlap was called when at least half of a TFBS overlapped with an L1PA2 transposon, using the –f option (Quinlan and Hall 2010). The number of TFBSs in each 100bp bin was counted using BedTools Coverage (Quinlan and Hall 2010), and normalised to the total counts of TFBSs mapped to the corresponding 20kb regions to obtain the normalised counts. The normalised counts in each bin were averaged across all L1PA2 elements containing at least one TFBS.

### Identifying L1PA2-derived TFBSs in MCF7 breast cancer cells

All TFBSs identified in MCF7 cells were obtained from the ChIP-Atlas database (Cell Type = “MCF-7”) (Oki et al. 2018). For the purpose of consistency, we referred to the TFs by the names recorded in ChIP-Atlas, instead of converting them to the protein names (e.g. ESR1 instead of ERα). The genomic coordinates of the TFBSs were converted from hg19 to hg38 using LiftOver, and only TFBSs that were successfully converted were retained for downstream analysis (Hinrichs et al. 2006). The genomic locations of the TFBSs were intersected with the L1PA2 elements using BedTools Intersect, where an overlap was called when at least half of a TFBS overlapped with an L1PA2 transposon, using the –f option (Quinlan and Hall 2010). The number of L1PA2 elements containing at least one TFBS in MCF7 cells was counted using a custom python script, and the lengths of L1PA2 transposons were calculated from their genomic coordinates. The number of L1PA2 overlaps for each TF and the number of TFs binding to each L1PA2 were counted using a custom python script.

### Epigenetic profiling of TF-bound L1PA2 transposons in MCF7 cells

L1PA2 transposons were divided into two groups based on the TF binding status in MCF7 cells. L1PA2 transposons were investigated for DNAse sensitivity, as well as active (H3K27ac, H3K4me1, H3K4me3 and H3K36me3) and repressive (H3K9me3 and H3K27me3) histone tail modifications in MCF7 cells. More specifically, epigenetic states of the 20kb region (bin = 100bp) centred on each L1PA2 transposon were investigated using publicly available DNAse-seq and ChIP-seq datasets, as described in Jiang and Upton (2019).

### Investigating TF co-localisation in L1PA2 transposons

To remove redundant information, overlapping L1PA2-derived binding sites of the same TF were merged to keep the most distal genomic coordinates. Locations of TFBSs in L1PA2 transposons were calculated as the distances between the middle of the binding sites and the 5’ ends of the transposons. The equivalent locations in the L1PA2 consensus sequence were then calculated by adjusting the distances by the RepStart and RepEnd information from the RepeatMasker database (Karolchik et al. 2004; Smit 2013-2015). To investigate TFBS co-localisation, for each pair of L1PA2-binding TFs, the number of L1PA2 transposons containing both TFBSs was counted and the distances between the binding sites were calculated using a custom python script.

### Co-localisation enrichment in L1PA2 transposons

Genome-wide binding sites for the top binding TFs (ESR1, SFPQ, MYC, FOXA1, NR2F2, CTCF, E2F1, KDM5B, ZNF143) were identified using the MCF7 ChIP-seq data from ChIP-Atlas (Oki et al. 2018). For each TF, non-L1PA2-derived TFBSs were identified by filtering out the previously identified L1PA2-derived TFBSs from genome-wide binding sites. Genomic locations of overlapping TFBSs were merged using BedTools Merge to remove redundant information (Quinlan and Hall 2010).

Here, we defined co-localised, or co-occurring TFBSs to be located within 500bp of each other. For each TF, binding sites that co-localised with another TF were identified using BedTools Window (`-w 500, -u`). The proportion of co-localisation was calculated as the number of co-localising TFBSs divided by the total number of TFBSs for this TF. To account for asymmetrical co-localisation, the co-localisation proportions for all TF-TF combinations were calculated in both orientations. For example, the proportion of ESR1 binding sites co-localising with SFPQ, and the proportion of SFPQ binding sites co-localising with ESR1, were calculated separately. The co-localisation proportions for L1PA2-derived and non-L1PA2-derived TFBSs were determined separately and compared using a one-tailed proportion z-test. To account for multiple testing, p-value 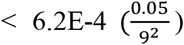 indicated statistical significance.

### Functional annotation of TF binding in L1PA2 transposons

To estimate the interactions between these TFs, the list of TFs binding to L1PA2 transposons in MCF7 cells was used as input for the online Multiple Proteins by Names / Identifiers search in STRING v11 (Szklarczyk et al. 2019). Only protein-protein interactions with experimental evidence were considered (active interaction sources = “Experiments”, minimum required interaction score = 0.15). In order to gain further insight into the pathways or diseases the TF networks were associated with, functional enrichment analysis was performed using the ToppFun tool (Chen et al. 2009). Bonferroni q-value < 0.05 indicated statistical significance.

### Motif analysis of TF binding in L1PA2 transposons

Binding motif position-weight matrices were obtained from the JASPAR 2020 database for ESR1 (MA0112.2), FOXA1 (MA0148.4) and E2F1 (MA0024.2) (Fornes et al. 2019). DNA sequences of L1PA2 transposons bound by each TF were extracted from the hg38 fasta file for main chromosomes (Lander et al. 2001; Kent et al. 2002) using BedTools Getfasta (v2.25.0) (`-s option`) (Quinlan and Hall 2010). Motif occurrences in L1PA2 sequences were identified using FIMO v5.0.5 (`-norc option`), with a statistical significance threshold of p-value < 1E-4 (Grant et al. 2011). Positions of the discovered motifs in the L1PA2 consensus sequence were calculated using the RepStart and RepEnd information from RepeatMasker (Smit 2013-2015). The distribution of discovered motifs in the consensus sequence (between 0 and 6500bp) was visualised by plotting the middle of the adjusted positions in a histogram (bin = 5bp). For bins that contained a high number of discovered motif sequences (peaks), a sequence logo, which was a representation of sequence conservation, was generated with the motif sequences in the corresponding bins using the WebLogo online tool v2.8.2 (Crooks et al. 2004). For the purpose of comparison, an L1PA2-specific sequence logo was generated for each TF using all discovered motif sequences.

### Identifying differentially expressed transcripts in MCF7 cells

SRA files of publicly available RNA-seq data in MCF7 and MCF10A cells (GSE108541, n = 3) were downloaded from the GEO database and converted to paired-end FASTQ files using command `fasterq-dump –S` (http://ncbi.github.io/sra-tools/) (SRA Toolkit Development Team) (Barrett et al. 2013). The quality of the sequencing data was confirmed using FastQC v0.11.8 (www.bioinformatics.babraham.ac.uk/projects/fastqc/) (Andrews 2010). Reads were aligned to GENCODE v29 transcripts (hg38) using STAR 2.7.1a, with more permissive multi-mappers parameters (`--outFilterMultimapNmax 100 --winAnchorMultimapNmax 200`, defaults are 20 and 50 respectively) (Dobin et al. 2013; Frankish et al. 2019). Transcripts were called and quantified for expression using StringTie 2.0.3 (`–rf -f 0.0 -c 0.001`) (Pertea et al. 2015). To include all annotated transcripts in our analysis, transcripts identified in each cell line were merged using StringTie 2.0.3 Merge (Pertea et al. 2015), and expression quantification was repeated.

Differential transcript expression analysis was performed using Ballgown 2.16.0 in R 3.6.1 (Frazee et al. 2015). Differentially expressed transcripts were defined as those with a fold change of more than 2 (up-regulated), or less than 0.5 (down-regulated) in MCF7 cells relative to MCF10A cells.

We defined the TSS of a transcript to be the position at which the transcript began. L1PA2-derived transcripts were identified by intersecting the TSSs of differentially expressed transcripts with L1PA2 transposons using BedTools Intersect. The exon structure and genomic locations of unknown transcripts were compared to nearby known transcripts by eye using the UCSC genome browser and information from the Ensembl database (Kent et al. 2002; Aken et al. 2016). Unknown transcripts were considered to be associated with a known transcript if they overlapped by at least one exon.

### Enrichment analysis of differentially expressed TSSs in L1PA2 transposons

L1PA2 transposons were split into two groups based on their TF binding status. For each group, we identified the L1PA2 transposons that were located within 1kb to 20kb (in increments of 1kb) away from an up-regulated or down-regulated TSS, using the BedTools Window -w option. The proportions of L1PA2 transposons located nearby the TSSs were calculated by dividing the counts by the total number of L1PA2 transposons in the corresponding TF-binding group.

To test for the statistical significance of direct overlaps between TSSs and L1PA2 transposons, the expected numbers of up-regulated and down-regulated TSSs in TF-bound L1PA2 transposons were estimated by random rotation of the genome and the TSS locations (10,000 permutations), using a custom python script. The average number of overlaps was divided by the total number of TSSs for up-regulated or down-regulated transcripts to produce the expected probability of TSSs being located in L1PA2 by random chance. The observed number of TSSs overlapping L1PA2 transposons was counted with no rotation applied, and the statistical significance for enrichment or depletion was evaluated using a binomial test. P-value < 0.05 indicated statistical significance.

### Epigenetic profiling of L1PA2-derived TSSs in MCF7 cells

L1PA2 transposons harbouring TSSs of up-regulated transcripts were investigated for their epigenetic states and RNA pol II binding. Epigenetic states of the 20kb region centred on each L1PA2 transposon (bin = 100bp) in MCF7 cells were investigated using publicly available ChIP-seq and DNAse-seq datasets, as described in Jiang and Upton (2019)). Similarly, the epigenetic profiles of the same regions in normal primary breast tissues were investigated (for data sources see Supplementary Table 4 and (Jiang and Upton 2019)).

RNA pol II binding data in breast tissues and cell lines (hg19) were downloaded from ChIP-Atlas (Oki et al. 2018) in the bed format (Cell Type Class = “Breast”), and converted to hg38 using LiftOver as previously described (Hinrichs et al. 2006). Overlapping binding sites were merged using BedTools Merge (Quinlan and Hall 2010). RNA pol II binding sites were intersected with the L1PA2 transposons using BedTools Intersect (Quinlan and Hall 2010), retaining information on the cell lines or tissues in which the binding sites were detected.

## DATA ACCESS

All publicly available datasets used in this study were referenced in the Methods section.

### ACKNOWLEDGMENTS

We thank the GTRD and ChIP-Atlas databases for compiling ChIP-seq data and making their results publicly available. K.U is supported by NHMRC Fellowship APP1130815. JC.J is supported by a UQ Research Training Scholarship.

## Author contributions

JC.J: conceptualisation, data analysis, data visualisation, manuscript writing, review and editing. J.R: supervision, manuscript review and editing. K.U: data curation, supervision, manuscript review and editing.

